# Cortical Sensitivity to Natural Scene Structure

**DOI:** 10.1101/613885

**Authors:** Daniel Kaiser, Greta Häberle, Radoslaw M. Cichy

## Abstract

Natural scenes are inherently structured, with meaningful objects appearing in predictable locations. Human vision is tuned to this structure: When scene structure is purposefully jumbled, perception is strongly impaired. Here, we tested how such perceptual effects are reflected in neural sensitivity to scene structure. During separate fMRI and EEG experiments, participants passively viewed scenes whose spatial structure (i.e., the position of scene parts) and categorical structure (i.e., the content of scene parts) could be intact or jumbled. Using multivariate decoding, we show that spatial (but not categorical) scene structure profoundly impacts on cortical processing: Scene-selective responses in occipital and parahippocampal cortices (fMRI) and after 255ms (EEG) accurately differentiated between spatially intact and jumbled scenes. Importantly, this differentiation was more pronounced for upright than for inverted scenes, indicating genuine sensitivity to spatial structure rather than sensitivity to low-level attributes. Our findings suggest that visual scene analysis is tightly linked to the spatial structure of our natural environments. This link between cortical processing and scene structure may be crucial for rapidly parsing naturalistic visual inputs.

## Cortical Sensitivity to Natural Scene Structure

Humans can efficiently extract information from natural scenes even from just a single glance (Potter, 1975; Thorpe et al., 1996). A major reason for this perceptual efficiency lies in the structure of natural scenes: for instance, a scene’s spatial structure tells us where specific objects can be found and its categorical structure tells us which objects are typically encountered within the scene (Kaiser et al., 2019; Oliva and Torralba, 2007; Võ et al., 2019; Wolfe et al., 2011).

The beneficial impact of scene structure on perception becomes apparent in jumbling paradigms, where the scene’s structure is purposefully disrupted by shuffling blocks of information across the scene. For instance, jumbling makes it harder to categorize scenes (Biederman et al., 1974), recognize objects within them (Biederman, 1972; Biederman et al., 1973) or to detect subtle visual changes (Varakin and Levin, 2008; Zimmermann et al., 2010). These findings suggest that typical scene structure contributes to efficiently perceiving a scene and its contents.

Such perceptual effects prompt the hypothesis that scene structure also impacts perceptual stages of cortical scene processing. However, while there is evidence that real-world structure impacts visual cortex responses to everyday objects (Kim and Biederman, 2011; Kaiser and Cichy, 2018; Kaiser and Peelen, 2018; Roberts and Humphreys, 2010) and human beings (Bernstein et al., 2010; Brandman and Yovel, 2016; Chan et al., 2010), it is unclear whether real-world structure has a similar impact on scene-selective neural responses.

To answer this question, we conducted multivariate pattern analysis (MVPA) and univariate analyses on fMRI and EEG responses to intact and jumbled scenes, which allowed us to spatially and temporally resolve whether cortical scene processing is indeed sensitive to scene structure. During the fMRI and EEG experiments, participants viewed scene images in which we manipulated two facets of natural scene structure: We orthogonally jumbled the scene’s spatial structure (i.e., whether the scene’s parts appear in their typical positions or not) or its categorical structure (i.e., whether the scene’s parts belong to the same category or different categories).

Our results provide three key insights into how scene structure affects scene representations: (1) Cortical scene processing is primarily sensitive to the scene’s spatial structure, more so than to the scene’s categorical structure. (2) Spatial structure impacts the perceptual analysis of scenes, in occipital and parahippocampal cortices (Epstein, 2012) and shortly after 200ms (Harel et al., 2016). (3) Spatial structure impacts cortical responses more strongly for upright than inverted scenes, indicating robust sensitivity to spatial scene structure that goes beyond sensitivity to low-level features.

## Materials and Methods

### Participants

In the fMRI experiment, 20 healthy adults participated in session 1 (mean age 25.5, *SD*=4.0; 13 female) and 20 in session 2 (mean age 25.4, *SD*=4.0; 12 female). Seventeen participants completed both sessions, three participants only session 1 or session 2, respectively. In the EEG experiment, 20 healthy adults (mean age 26.6, *SD*=5.8; 9 female) participated in a single session. Samples sizes were determined based on typical samples sizes in related research; a sample of *N*=20 yields 80% power for detecting effects sizes greater than *d*=0.66^1^. All participants had normal or corrected-to-normal vision. Participants provided informed consent and received monetary reimbursement or course credits. All procedures were approved by the ethical committee of Freie Universität Berlin and were in accordance with the Declaration of Helsinki.

### Stimuli and design

Stimuli were 24 scenes from four different categories (church, house, road, supermarket; Figure 1a), taken from an online resource (Konkle et al., 2010); the complete scene image set can be found in the Supplementary Information. We split each image into quadrants and systematically recombined the resulting parts in a 2×2 design, where both the scenes’ spatial structure and their categorical structure could be either intact or jumbled (Figure 1b/c). This yielded four conditions: (1) In the “spatially intact & categorically intact” condition, parts from four scenes of the same category were combined in their correct locations. (2) In the “spatially intact & categorically jumbled” condition, parts from four scenes from different categories were combined in their correct locations. (3) In the “spatially jumbled & categorically intact” condition, parts from four scenes of the same category were combined, and their locations were exchanged in a crisscrossed way. (4) In the “spatially jumbled & categorically jumbled” condition, parts from four scenes from different categories were combined, and their locations were exchanged in a crisscrossed way. For each participant separately, 24 unique stimuli were generated for each condition by randomly drawing suitable fragments from different scenes^2^. During the experiment, all scenes were presented both upright and inverted.

**Figure 1.**
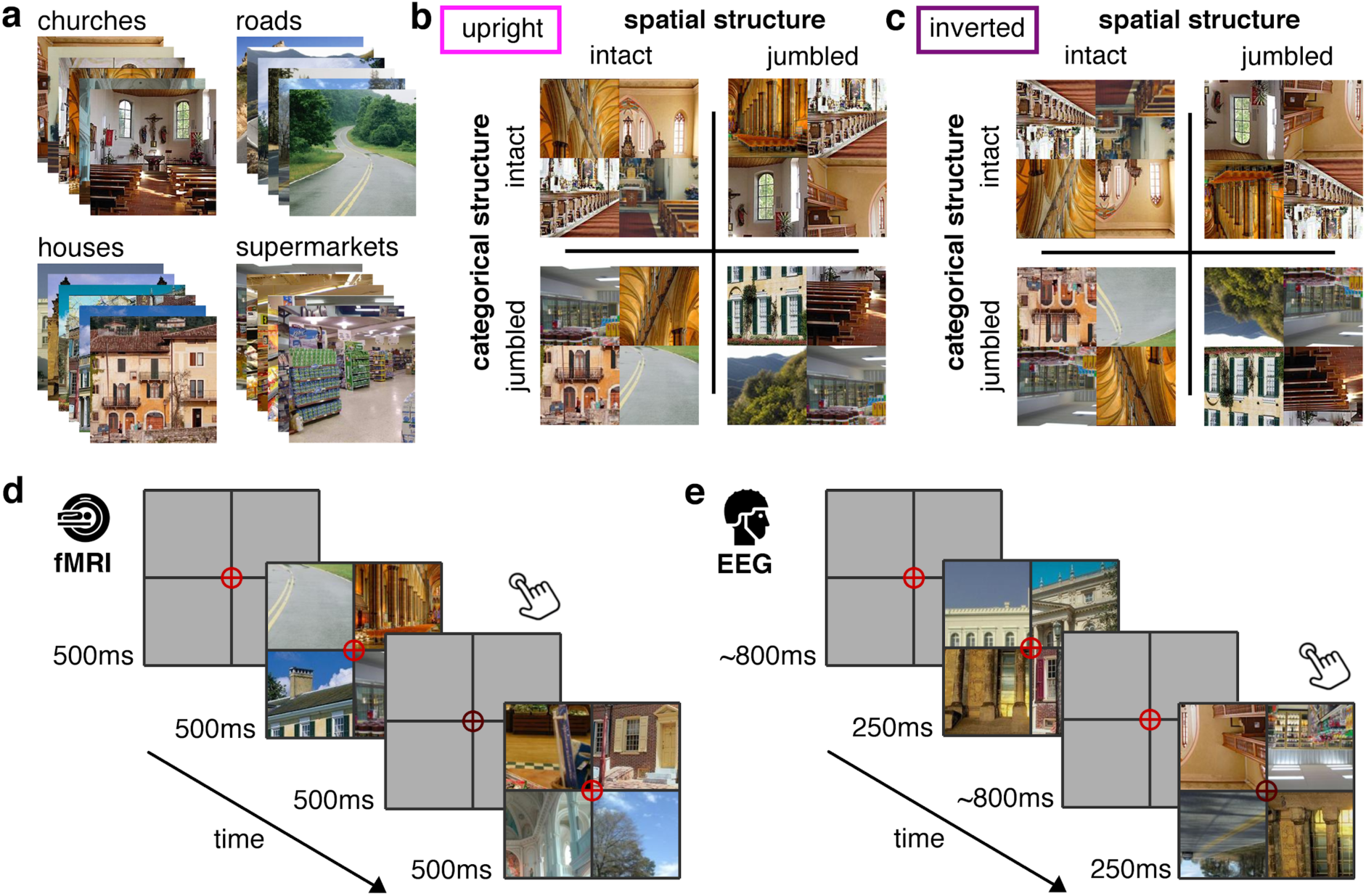
Stimuli and Paradigm. We combined parts from 24 scene images from four categories (a) to create a stimulus set where the scenes’ structural (e.g. the spatial arrangements of the parts) and their categorical structure (e.g., the category of the parts) was orthogonally manipulated; all scenes were presented both upright and inverted (b/c). In the fMRI experiment, scenes were presented in a block design, where each block of 24s exclusively contained scenes of a single condition (d). In the EEG experiment, all conditions were randomly intermixed (e). During both experiments, participants responded to color changes of the central crosshair.

### fMRI paradigm

The fMRI experiment (Figure 1d) comprised two sessions. In the first session, upright scenes were shown, in the second session inverted scenes were shown; the sessions were otherwise identical. Each session consisted of five runs of 10min. Each run consisted of 25 blocks of 24 seconds. In 20 blocks, scene stimuli were shown with a frequency of 1Hz (0.5s stimulus, 0.5s blank). Each block contained all 24 stimuli of a single condition. In 5 additional fixation-only blocks, no scenes were shown. Block order was randomized within every five consecutive blocks, which contained each condition (four scene conditions and fixation-only) exactly once.

Scene stimuli appeared in a black grid (4.5° visual angle), which served to mask visual discontinuities between quadrants. Participants were monitoring a central red crosshair, which twice per block (at random times) darkened for 50ms; participants had to press a button when they detected a change. Participants on average detected 80.0% (*SE*=2.5)^3^ of the changes. Stimulus presentation was controlled using the Psychtoolbox (Brainard, 1997).

In addition to the experimental runs, each participant completed a functional localizer run of 13min, during which they viewed images of scenes, objects, and scrambled scenes. The scenes were new exemplars of the four scene categories used in the experimental runs; objects were also selected from four categories (car, jacket, lamp, sandwich). Participants completed 32 blocks (24 scene/object/scrambled blocks and 8 fixation-only blocks), with parameters identical to the experimental runs (24s block duration, 1Hz stimulation frequency, color change task).

### EEG paradigm

In the EEG experiment (Figure 1e), all conditions were randomly intermixed within a single session of 75min (split into 16 runs). During each trial, a scene appeared for 250ms, followed by an inter-trial interval randomly varying between 700ms and 900ms. In total, there were 3072 trials (384 per condition), and an additional 1152 target trials (see below).

As in the fMRI, stimuli appeared in a black grid (4.5° visual angle) with a central red crosshair. In target trials, the crosshair darkened during the scene presentation; participants had to press a button and blink when detecting this change. Participants on average detected 78.1% (*SE*=3.6) of the changes. Target trials were not included in subsequent analyses.

### fMRI recording and preprocessing

MRI data was acquired using a 3T Siemens Tim Trio Scanner equipped with a 12-channel head coil. T2*-weighted gradient-echo echo-planar images were collected as functional volumes (*TR*=2s, *TE*=30ms, 70° flip angle, 3mm^3^ voxel size, 37 slices, 20% gap, 192mm FOV, 64×64 matrix size, interleaved acquisition). Additionally, a T1-weighted anatomical image (MPRAGE; 1mm^3^ voxel size) was obtained. Preprocessing was performed using SPM12 (www.fil.ion.ucl.ac.uk/spm/). Functional volumes were realigned, coregistered to the anatomical image, and normalized into MNI-305 space. Images from the localizer run were additionally smoothed using a 6mm full-width-half-maximum Gaussian kernel.

### EEG recording and preprocessing

EEG signals were recorded using an EASYCAP 64-electrode^4^ system and a Brainvision actiCHamp amplifier. Electrodes were arranged in accordance with the 10-10 system. EEG data was recorded at 1000Hz sampling rate and filtered online between 0.03Hz and 100Hz. All electrodes were referenced online to the Fz electrode. Offline preprocessing was performed using FieldTrip (Oostenveld et al., 2011). EEG data were epoched from −200ms to 800ms relative to stimulus onset, and baseline-corrected by subtracting the mean pre-stimulus signal. Channels and trials containing excessive noise were removed based on visual inspection. Blinks and eye movement artifacts were removed using independent component analysis and visual inspection of the resulting components. The epoched data were down-sampled to 200Hz.

### fMRI region of interest definition

We restricted fMRI analyses to three regions of interest (ROIs): early visual cortex (V1), scene-selective occipital place area (OPA), and scene-selective parahippocampal place area (PPA) (Figure 2). We additionally localized scene-selective retrosplenial cortex (RSC), but did not observe reliable above-baseline activations to our scene stimuli in this region, all *t*(19)<0.14, *p*>.45. The results for RSC can be found in the Supplementary Information.

**Figure 2.**
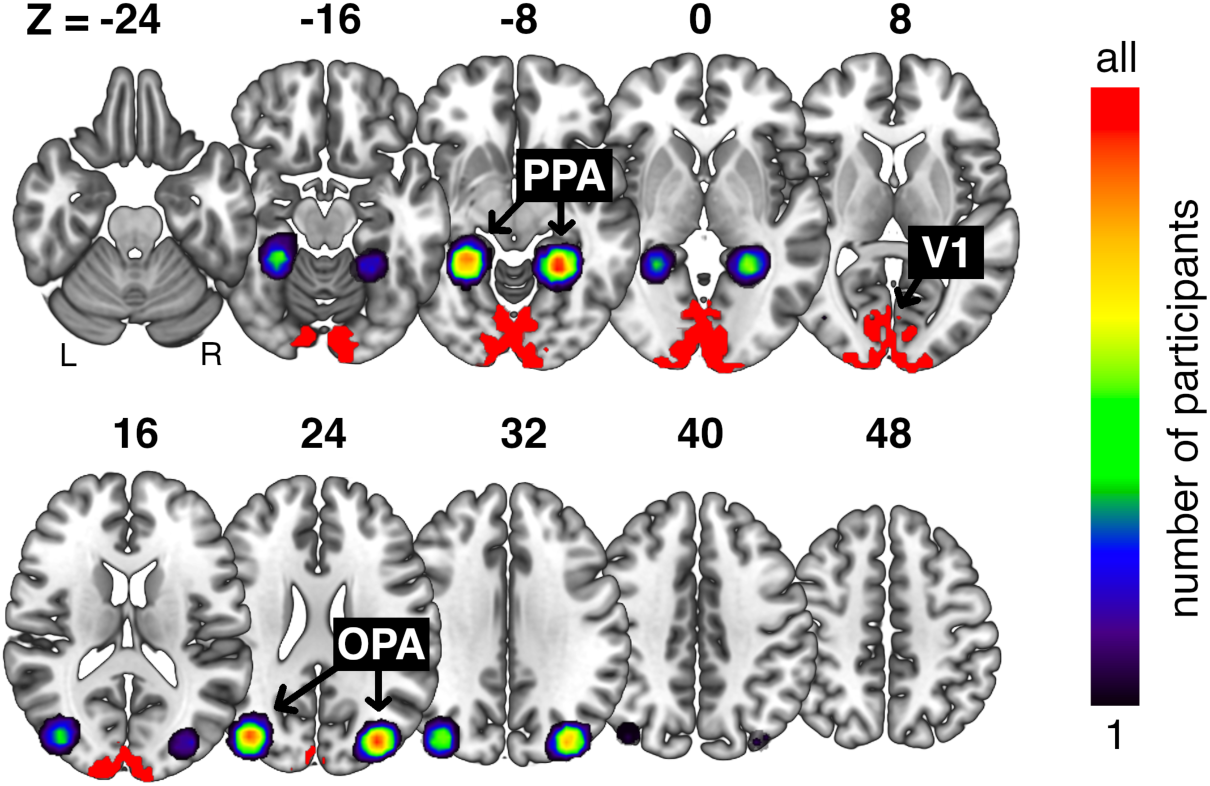
Location of the fMRI regions of interest (ROIs). fMRI data analysis was restricted to three ROIs: primary visual cortex (V1), the occipital place area (OPA) and the parahippocampal place area (PPA). The V1 ROI was based on a functional atlas (Wang et al., 2015), and identical for all participants. The scenes-selective regions were defined as spheres around each participant’s peak activation in a separate scene-localizer run, constrained by functional group masks (Julian et al., 2012). The colormap represents the consistency of ROI locations across participants (i.e., how many participants’ ROIs covered the respective voxels).

V1 was defined based on a functional group atlas (Wang et al., 2015), from which we selected all voxels that had a higher probability of belonging to V1 than belonging to another region in the atlas (905 voxels). Changing the number of voxels included did not qualitatively change the results in V1 (see Supplementary Information).

Scene-selective ROIs were defined using the localizer data, which were modelled in a general linear model (GLM) with 9 predictors (3 regressors for the scene/object/scrambled blocks and 6 movement regressors). Scene-selective ROI definition was constrained by group-level activation masks for OPA and PPA (Julian et al., 2012). Within these masks, we first identified the voxel exhibiting the greatest *t*-value in a scene>object contrast, separately for each hemisphere, and then defined the ROI as a 125-voxel sphere around this voxel (similar results were obtained for different ROI sizes, see Supplementary Information). Left- and right-hemispheric ROIs were concatenated for further analysis^5^.

### fMRI decoding

fMRI response patterns for each ROI were extracted directly from the volumes recorded during each block. After shifting the activation time course by three TRs (i.e., 6s) to account for the hemodynamic delay, we extracted voxel-wise activation values from the 12 TRs corresponding to each block of 24s. Activation values for these 12 TRs were then averaged, yielding a single response pattern across voxels for each block. To account for activation differences between runs, the mean activation across all blocks was subtracted from each voxel’s values, separately for each run. Decoding analyses were performed using CoSMoMVPA (Oosterhof et al., 2016), and were carried out separately for each ROI and participant. We used data from four runs to train linear discriminant analysis (LDA) classifiers to discriminate multi-voxel response patterns (i.e., patterns of voxel activations across all voxels of an ROI) for two conditions (e.g., spatially intact versus spatially jumbled scenes). Classifiers were tested using response patterns for the same two conditions from the left-out, fifth run. This classification routine was done repeatedly until every run was left out once and decoding accuracy was averaged across these repetitions.

### fMRI univariate analysis

To establish univariate activation differences, we modeled the fMRI data in a GLM analysis. For this analysis, all functional volumes were smoothed using a 6mm full-width-half-maximum Gaussian kernel. For each run, we constructed a GLM with 10 predictors (4 regressors reflecting the four scene conditions and 6 movement regressors). For each of the four scene conditions, this analysis yielded 5 beta maps (one for each run) for the upright scenes (from session 1), and 5 beta maps (one for each run) for the inverted scenes (from session 2). We first averaged beta weights for every condition across runs. These beta weights were then averaged across all voxels of each ROI, yielding one activation value for each condition, ROI, and participant. For each ROI (V1, OPA, PPA), and separately for the two stimulus orientations (upright, inverted), we computed three effects: (1) The main effect of spatial structure, reflecting the difference between the two spatially intact and the two spatially jumbled scenes, (2) the main effect of categorical structure, reflecting the difference between the two categorically intact and the two categorically jumbled scenes, and (3) the interaction effect of spatial and categorical structure. Subsequently, to uncover inversion effects, we compared these effects across the upright scenes and inverted scenes to reveal inversion effects.

### EEG decoding

EEG decoding was performed separately for each time point (i.e., every 5ms) from −200ms to 800ms relative to stimulus onset, using CoSMoMVPA (Oosterhof et al., 2016). We used data from all-but-one trials for two conditions to train LDA classifiers to discriminate topographical response patterns (i.e., patterns across all electrodes) for two conditions (e.g., spatially intact versus spatially jumbled scenes). Classifiers were tested using response patterns for the same two conditions from the left-out trials. This classification routine was done repeatedly until each trial was left out once and decoding accuracy was averaged across these repetitions. Classification time series for individual participants were smoothed using a running average of five time points (i.e., 25ms).

### EEG univariate analysis

To establish univariate EEG response differences (i.e., ERP effects) between conditions, we averaged evoked responses for all trials of each condition. Based on a previous study on scene-selective ERPs (Harel et al., 2016), we then averaged these responses across six posterior-lateral EEG electrodes (P4, P8, O2, P7, P3, O1), yielding one ERP response for each condition and participant. For these ERPs, we computed the same effects as outlined above for the fMRI data: a main effect of spatial structure, a main effect of categorical structure, and interactions with scene inversion^6^.

### Statistical testing

For the fMRI data, we used *t*-tests to compare decoding against chance and between conditions. For the univariate data, we used ANOVAs to tests for differences in activations. To Bonferroni-correct for comparisons across ROIs, all *p*-values were multiplied by 3. For the EEG data, given the larger number of comparisons, we used a threshold-free cluster enhancement procedure (Smith and Nichols, 2009) and multiple-comparison correction based on a sign-permutation test (with null distributions created from 10,000 bootstrapping iterations), as implemented in CoSMoMVPA (Oosterhof et al., 2016). The resulting statistical maps were thresholded at *z*>1.96 (i.e., *p*_*corr*_<.05).

### Data Availability

Data are publicly available on OSF (doi.org/10.17605/OSF.IO/W9874). Materials and code are available from the corresponding author upon request.

## Results

For both the fMRI and EEG data, we performed two complimentary decoding analyses. In the first analysis, we tested sensitivity for spatial structure by decoding spatially intact from spatially jumbled scenes (Figure 3a). In the second analysis, we tested sensitivity for categorical structure by decoding categorically intact from categorically jumbled scenes (Figure 3d). To investigate whether successful decoding indeed reflected sensitivity to scene structure, we performed both analyses separately for the upright and inverted scenes. Critically, inversion effects (i.e., better decoding in the upright than in the inverted condition) indicate genuine sensitivity to natural scene structure that goes beyond purely visual differences.

**Figure 3.**
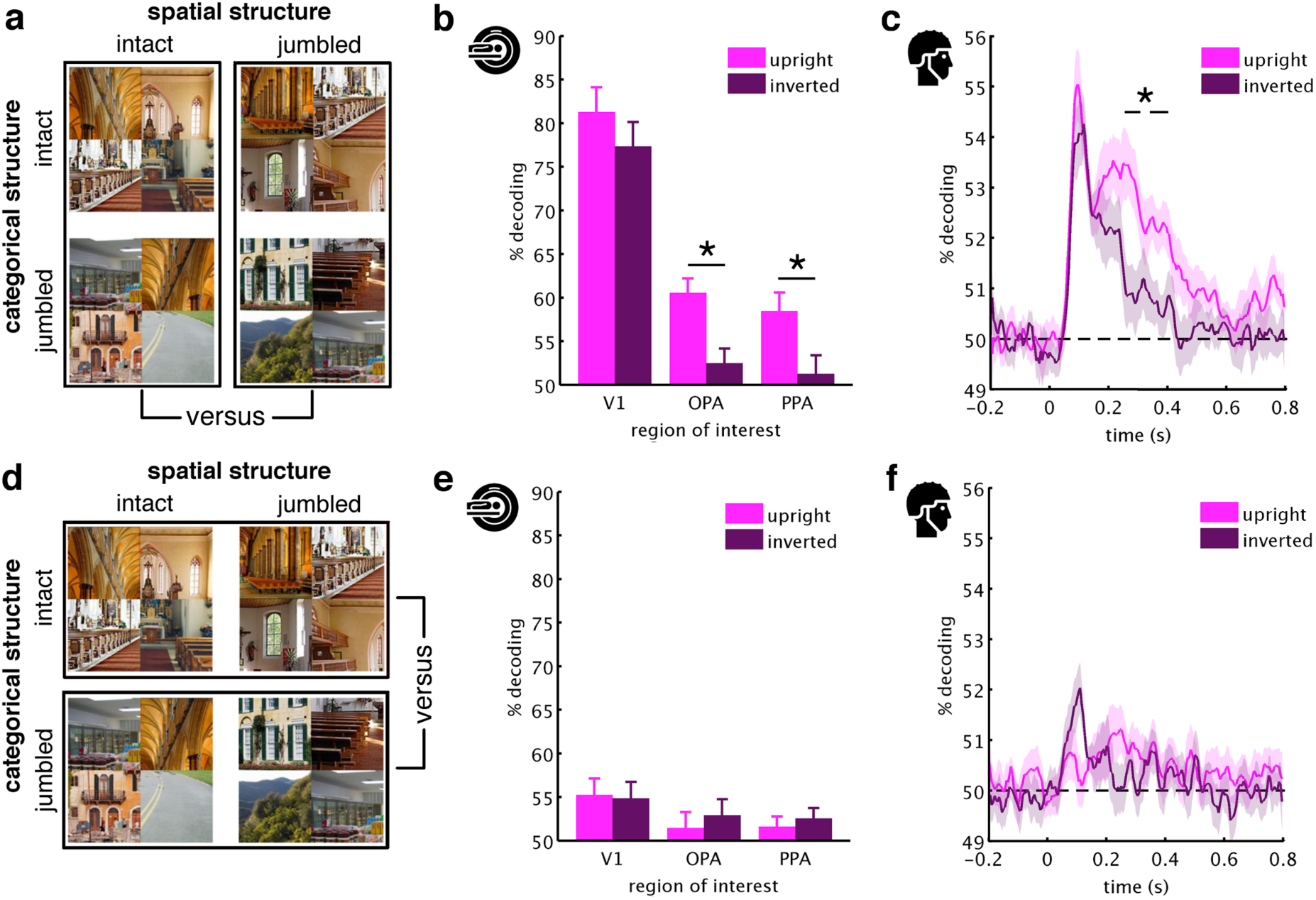
MVPA results. To reveal sensitivity to spatial scene structure, we decoded between scenes with spatially intact and spatially jumbled parts (a). Already during early processing (in V1 and before 200ms) spatially intact and jumbled scenes could be discriminated well, both for the upright and inverted conditions. Critically, during later processing (in OPA/PPA and from 255ms) inversion effects (i.e., better decoding for upright than inverted scenes) revealed genuine sensitivity to spatial scene structure (b/c). To reveal sensitivity to categorical scene structure, we decoded between scenes with categorically intact and categorically jumbled parts (d). In this analysis, no pronounced decoding and no inversion effects were found, neither across space (e) nor time (f). Error margins reflect standard errors of the difference. Significance markers denote inversion effects (*p*_*corr*_<.05).

### Sensitivity to spatial scene structure

First, to uncover where and when cortical processing is sensitive to spatial structure, we decoded between scenes whose spatial structure was intact or jumbled (Figure 3a).

For the fMRI data (Figure 3b), we found highly significant decoding between spatially intact and spatially jumbled scenes. For upright scenes, significant decoding emerged in V1, *t*(19)=13.03, *p*_*corr*_<.001, OPA, *t*(19)=7.61, *p*_*corr*_<.001, and PPA, *t*(19)=5.92, *p*_*corr*_=.002, and for inverted scenes in V1, *t*(19)=9.92, *p*_*corr*_<.001, but not in OPA, *t*(19)=2.08, *p*_*corr*_=.16, and PPA, *t*(19)=0.85, *p*_*corr*_>1. Critically, we observed inversion effects (i.e., better decoding for the upright scenes) in the OPA, *t*(16)=4.41, *p*_*corr*_=.001^7^, and PPA, *t*(16)=3.67, *p*_*corr*_=.006, but not in V1, *t*(16)=1.32, *p*_*corr*_=.62. Therefore, decoding in V1 solely reflects visual differences, whereas OPA and PPA exhibit genuine sensitivity to the spatial scene structure. This result was confirmed by further ROI analyses and a spatially unconstrained searchlight analysis (see Supplementary Information).

For the EEG data (Figure 3c), we also found strong decoding between spatially intact and jumbled scenes. For upright scenes, this decoding emerged between 55ms and 465ms, between 505ms and 565ms, and between 740ms and 785ms, peak *z*>3.29, *p*_*corr*_<.001, and for inverted scenes between 65ms and 245ms, peak *z*>3.29, *p*_*corr*_<.001. As in scene-selective cortex, we observed inversion effects, indexing stronger sensitivity to spatial structure in upright scenes, between 255ms and 300ms and between 340ms and 395ms, peak *z*=2.78, *p*_*corr*_=.005.

Together, these results show that in scene-selective OPA and PPA, and after 255ms, cortical activations are sensitive to the spatial structure of natural scenes. Critically, this sensitivity becomes apparent in inversion effects, and thus cannot be attributed to image-specific differences between intact and jumbled scenes, as these are identical for the upright and inverted scenes. Our findings rather indicate a genuine sensitivity to spatial structure consistent with real-world experience.

### Sensitivity to categorical scene structure

Second, to uncover where and when cortical processing is sensitive to categorical structure, we decoded between scenes whose categorical structure was intact or jumbled (Figure 3a).

For the fMRI (Figure 3e), the upright scenes’ categorical structure could be decoded only from V1, *t*(19)=3.11, *p*_*corr*_=.017, but not the scene-selective ROIs, both *t*(19)<2.15, *p*_*corr*_>.13. Similarly, for the inverted scenes, significant decoding was only observed in V1, *t*(19)=4.58, *p*_*corr*_<0.001, but not in the scene-selective ROIs, both *t*(19)<2.29, *p*_*corr*_>.10. No inversion effects were observed, all *t*(16)<0.60, *p*_*corr*_>1.

For the EEG (Figure 3f), we found only weak decoding between the categorically intact and jumbled scenes. In the upright condition, decoding was significant between 165ms and 175ms and between 215ms and 265ms, peak *z*=2.32, *p*_*corr*_=.02, and in the inverted condition at 120ms, peak *z*=1.97, *p*_*corr*_=.049. No significant inversion effects were observed, peak *z*=1.64, *p*_*corr*_=.10^8^.

Together, these results reveal no substantial sensitivity to the categorical structure of a scene, at least when none of the scenes are fully coherent and when they are not relevant for behavior. Please note that this absence of an effect does not in no way entail that there is no representation of category during scene analysis. In our analysis, we did not decode between different scene categories, but between scenes whose categories were intact or shuffled (collapsed across their categorical content): as a consequence, our analysis only reveals an absence of sensitivity for categorical structure, but not an absence of sensitivity for category per se.

This absence of sensitivity for categorical scene structure is in marked contrast with sensitivity for spatial scene structure, which is observed in the absence of behavioral relevance and is disrupted by stimulus inversion.

### Enhanced responses to spatially structured scenes

Our decoding analyses show that scene-selective cortex exhibits a profound sensitivity to spatial scene structure. To further understand this sensitivity, we conducted a univariate analysis, in which we compared the magnitude of responses evoked by intact and jumbled scenes (Figure 4a/c). Critically, this analysis allowed us to disentangle two opposing interpretations: On one side, sensitivity to scene structure could indeed reflect a visual tuning to real-world properties – in this case, enhanced responses to intact scenes, compared to jumbled scenes, are expected. On the other side, sensitivity to scene structure could mainly reflect the coding of stimuli that are incoherent with real-world experience, reflecting a type of “surprise” response – in this case, enhanced responses to jumbled scenes, compared to intact scenes, are expected. Analyzing response magnitudes across space (fMRI) and time (EEG) allowed us to arbitrate these two interpretations.

**Figure 4.**
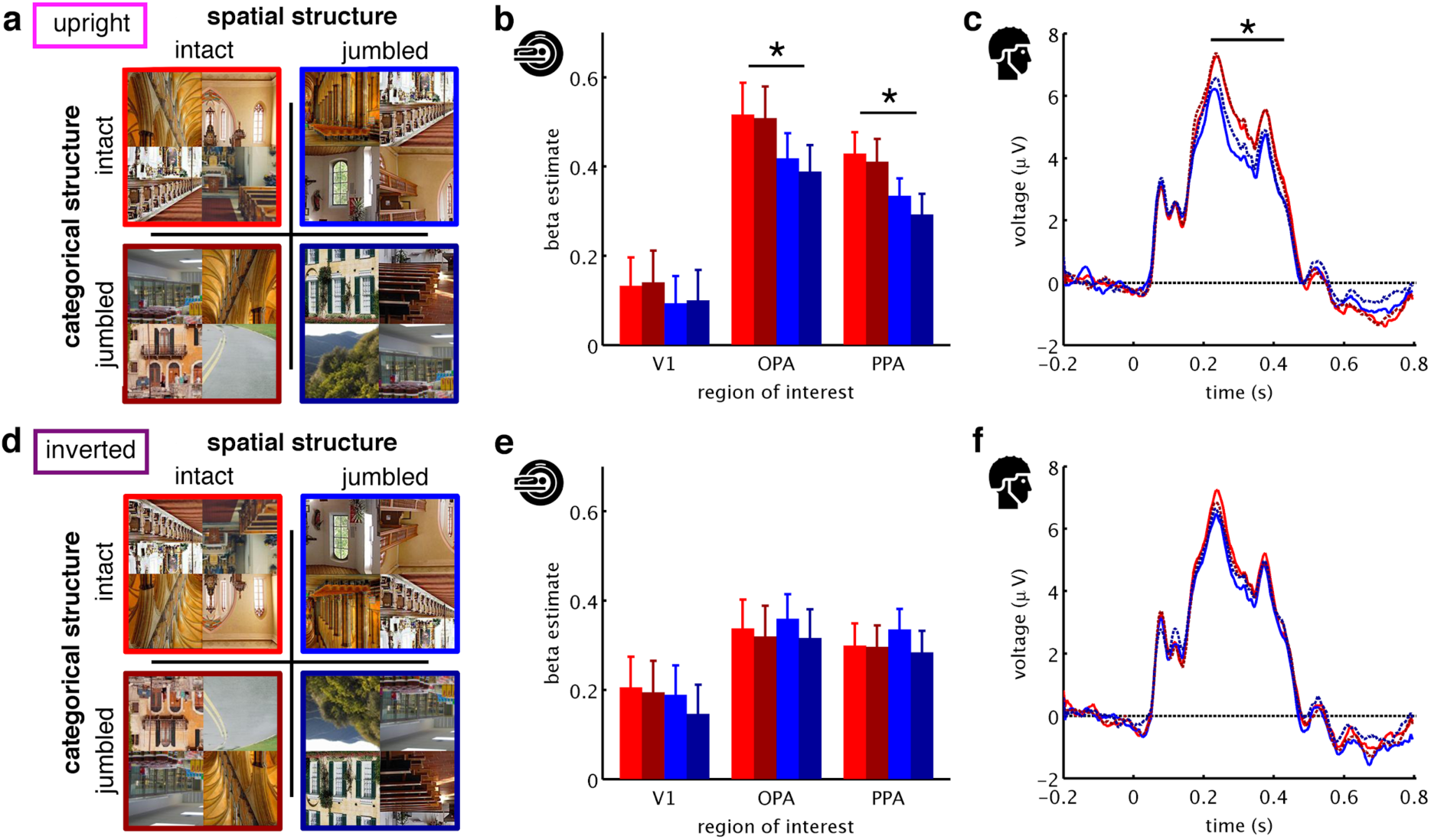
Univariate Results. To reveal sensitivity to scene structure in univariate response magnitudes, we looked at average responses to each of the four conditions, separately for the upright scenes (a) and the inverted scenes (d). For the upright scenes, we found main effects of spatial structure in OPA and PPA (b) and between 225ms and 425ms (c), while no effects of spatial structure were found for the inverted scenes (e/f). Supporting our MVPA results, inversion effects (i.e., greater effects of spatial structure in the upright, compared to the inverted scenes) were found in OPA and PPA (at *p*_*corr*_<.05) and from 260ms (at a more liberal *p*_*corr*_<.1), indicating increased responsiveness to spatially structured scenes. No main effects of categorical structure and no interaction effects were found. Error margins reflect standard errors of the mean. Significance markers denote main effects of spatial structure (*p*_*corr*_<.05).

In the fMRI, we found significant main effects of spatial structure in the upright condition in OPA, *F*(1,19)=21.00, *p*_*corr*_<.001, and PPA, *F*(1,19)=55.30, *p*_*corr*_<.001, but not in V1, *F*(1,19)=5.11, *p*_*corr*_=.11 (Figure 4b). No main effects of categorical structure, all *F*(1,19)<5.69, *p*_*corr*_>.08, and no interactions between spatial and categorical structure were found, all *F*(1,19)<1.18, *p*_*corr*_>.88. In the inverted condition, we observed no significant effects, all *F*(1,19)<1.12, *p*_*corr*_>.92 (Figure 4e). Critically, we inversion effects revealed greater effects of spatial structure in the upright than in the inverted condition in OPA, *F*(1,16)=17.04, *p*_*corr*_=.002, and PPA, *F*(1,16)=21.82, *p*_*corr*_<.001. In accordance with the MVPA results, this finding indicates genuine sensitivity to spatial scene structure in OPA and PPA. Additionally, the univariate results highlight that scene-selective cortex preferentially responds to the spatially intact scenes, rather than the spatially jumbled scenes.

In the EEG, we only found a significant main effect of spatial structure for the upright scenes (Figure 4c/f), which emerged between 225ms and 425ms, peak *z*=3.09, *p*_*corr*_=.002. None of the other main effects or interactions were significant. However, we observed trending inversion effects (at a more liberal threshold of *p*_*corr*_<.1), which emerged between 260ms and 270ms, and at 305ms, peak *z*=1.72, *p*_*corr*_=.086. Although not significant, these trending effects qualitatively resemble the findings obtained in the more sensitive MVPA, which showed that from 255ms responses become sensitive to spatial scene structure.

Together, the univariate results highlight that responses to natural scenes are stronger for scenes that are spatially structured. This suggests a preferential processing of scenes that are composed in accordance with real-world experience – rather than an enhanced response to scenes that do not adhere to this experience.

## Discussion

Our findings provide the first spatiotemporal characterization of cortical sensitivity to natural scene structure. As the key result, we observed sensitivity to spatial (but not categorical) scene structure, which emerged in scene-selective cortex and from 255ms of vision. By showing that this effect is stronger for upright than for inverted scenes, we provide strong evidence for genuine sensitivity to spatial structure, rather than low-level properties.

Sensitivity to spatial structure may index mechanisms enabling efficient scene understanding. Previous work on object processing shows that in order to efficiently parse the many objects contained in natural scenes, the visual system exploits regularities in the environment, such as regularities in individual objects’ positions (Kaiser and Cichy, 2018; Kaiser et al., 2018), relationships between objects (Kim and Biederman, 2011; Kaiser and Peelen, 2018; Kaiser et al., 2014; Roberts and Humphreys, 2010), and relationships between objects and scenes (Brandman and Peelen, 2017; Faivre et al., 2019). Further, a recent fMRI study suggests that low-level representations of small and incomplete scene fragments partly depend on the fragment’s typical position within the visual world (Mannion, 2015). Relatedly, we recently showed that in scene-selective occipital cortex and after 200ms of vision, the representations of such scene fragments are sorted with respect to their typical location in the world (Kaiser et al., 2019a). Focusing on the interplay of multiple scene elements, the current study shows that on higher levels of the scene processing hierarchy, the visual system uses spatial regularities to concurrently process the multiple elements of complex scenes in an efficient way. This result is in line with the emerging view that real-world structure facilitates processing in the visual system across diverse naturalistic contents (Kaiser et al., 2019b).

What mechanism underlies the preferential processing of spatially structured scenes? As one possibility, a scene’s intact spatial structure may trigger integrative processing across the scene, akin to integrative processing of multiple objects that are positioned in accordance with spatial regularities (Baldassano et al., 2017; Kaiser and Peelen, 2018). Alternatively, spatially structured scenes may contain typical global properties (Oliva and Torralba, 2006) that are absent in spatially jumbled scenes, and the sensitivity to spatial structure may partly reflect sensitivity to the formation of such global properties. At this point, more studies are needed to understand which types of features drive the sensitivity to spatial structure.

Our results also shine new light on the temporal processing cascade during scene perception. Sensitivity to spatial structure emerged after 255ms of processing, which is only after scene-selective peaks in ERPs (Harel et al., 2016; Sato et al., 1999)^9^ and after basic scene attributes are computed (Cichy et al., 2017). Interestingly, after 250ms brain responses not only become sensitive to scene structure, but also to object-scene consistencies (Draschkow et al., 2018; Ganis and Kutas, 2003; Mudrik et al., 2010; Võ and Wolfe, 2013). Together, these results suggest a dedicated processing stage for the structural analysis of objects, scenes, and their relationships, which is different from basic perceptual processing. However, whether these different findings indeed reflect a common underlying mechanism requires further investigation. For instance, future investigations need to clarify which of these studies effects reflect enhanced processing of consistent structure (as our study does) and which primarily reflect responses to inconsistencies.

Our findings suggest more pronounced sensitivity to spatial structure than to categorical structure. This is in line with studies showing that scene-selective responses are mainly driven by spatial layout, rather than scene content (Dillon et al., 2018; Harel et al., 2013; Henriksson et al., 2019; Kravitz et al., 2011). However, our results need not to be taken as evidence that categorical structure is not represented at all during visual analysis^10^. It is conceivable that visual processing is less sensitive to categorical structure when, as in our study, all scenes are jumbled to some extent and not behaviorally relevant.

On the contrary, robust sensitivity to spatial scene structure emerged in the absence of behavioral relevance. This suggests that spatial structure is analyzed automatically during perceptual processing and is not strongly dependent on attentional engagement with the scene. As in real-world situations we cannot explicitly engage with all aspects of a scene concurrently, this automatic analysis of spatial structure may be crucial for rapid scene understanding.

## Acknowledgements

We thank Sina Schwarze for help in EEG data collection and manuscript preparation. D.K. and R.M.C. are supported by Deutsche Forschungsgemeinschaft (DFG) grants (KA4683/2-1, CI241/1-1, CI241/3-1). R.M.C. is supported by a European Research Council Starting Grant (ERC-2018-StG 803370). The authors declare no conflict of interest.

## Author Contributions

D. K. and R.M.C. designed research, D.K. and G.H. acquired data, D.K. and G.H. analyzed data, D.K., G.H., and R.M.C. interpreted results, D.K. prepared figures, D.K. drafted manuscript, D.K., G.H., and R.M.C. edited and revised manuscript. All authors approved the final version of the manuscript.

## Footnotes

1 Related studies on object-object and object-scene consistencies typically yield large effect sizes which exceed this value, both for fMRI responses, *d*=0.72 (Brandman and Peelen, 2017), *d*=0.67 (Kaiser and Peelen, 2018), *d*=2.14 (Kim and Biederman, 2011), *d*=0.94 (Roberts and Humphreys, 2010), and EEG responses, *d*=0.71 (Draschkow et al., 2018), *d*=0.88 (Ganis and Kutas, 2003), *d*=0.67 (Mudrik et al., 2010), *d*=0.69 (Võ and Wolfe, 2013).

2 Note that all scenes were jumbled to some extent, as also in the categorically intact scenes four different exemplars were intermixed.

3 For two participants, due to technical problems, no button presses were recorded.

4 For two participants, due to technical problems, only data from 32 electrodes was recorded.

5 Analysing the data from the two hemispheres separately did not yield any significant differences between hemispheres (*F*<2.04, *p*>.17, for all interactions with hemisphere).

6 For using the same statistical tests as for the decoding results, interactions in the univariate EEG analyses were computed by testing the differences between conditions against each other (e.g., the difference between intact and jumbled scenes in the upright condition versus the difference between intact and jumbled scenes in the inverted conditions).

7 Statistics for fMRI inversion effects are based on the 17 participants who completed both sessions.

8 Note that the strongest tendency towards an inversion effect (at 115ms) was against the predicted direction.

9 In our study, ERP responses in posterior-lateral electrodes peaked at 235ms.

10 In the Supplementary Information, we show that the four scene categories can be successfully decoded from the EEG signals.

