## Supplementary Information for "Cortical Sensitivity to Natural Scene Structure"

**Contents:**

- Complete scene image set
- fMRI decoding – searchlight analysis
- fMRI decoding – additional ROIs
- fMRI decoding – varying voxel counts in V1
- fMRI decoding – varying voxel counts in OPA / PPA
- EEG decoding – classifying scene category

**Complete scene image set**

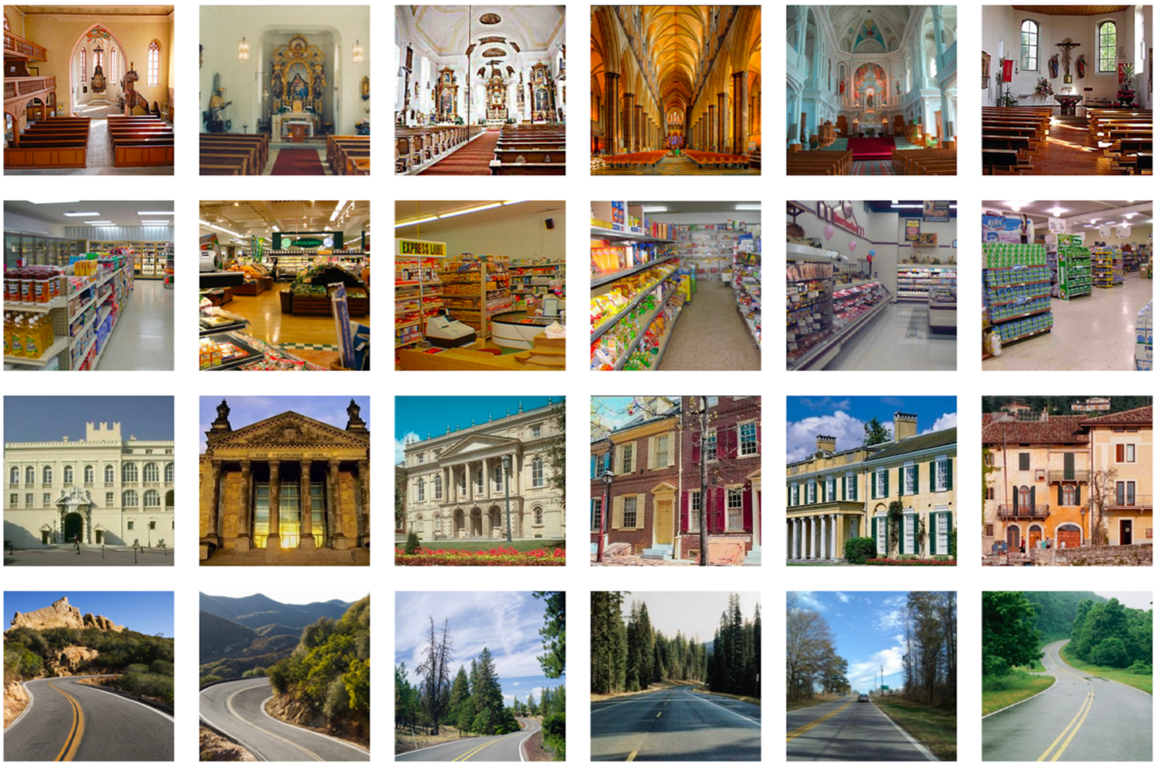

*Figure S1.* Scene images used in the study. The intact and jumbled scene stimuli were generated from 24 scene images from four categories: churches, supermarkets, houses, and streets. All images were chosen to depict easily recognizable scenes, photographed from a typical real-life viewpoint.

**fMRI decoding – searchlight analysis**

To substantiate our ROI analyses, we additionally ran a searchlight MVPA, where we probed sensitivity to spatial and categorical scene structure across the whole occipitotemporal visual cortex.

For this searchlight MVPA, we repeatedly performed the two decoding analyses (i.e., decoding spatial or categorical scene structure; see Method) for a moving sphere of 250 voxels, which was centered on every voxel within an anatomical mask of the occipital and temporal cortices (taken from WFU PickAtlas for SPM12). This procedure allowed us to map sensitivity to spatial and categorical scene structure across the whole visual cortex, separately for the upright and inverted scenes, and separately for each participant. By testing decoding against chance across participants, we computed six effects: (1) sensitivity to spatial structure for the upright scenes, (2) sensitivity to spatial structure for the inverted scenes, (3) sensitivity to categorical structure for the upright scenes, (4) sensitivity to categorical structure for the inverted scenes, (5) an inversion effect for spatial structure (i.e., the difference between (1) and (2)), and (6) an inversion effect for categorical structure (i.e., the difference between (3) and (4)). Significance was established using a threshold-free cluster enhancement procedure (as used for the EEG data). The resulting statistical maps were thresholded at $z>1.96$ (i.e., $p_{corr}<.05$).

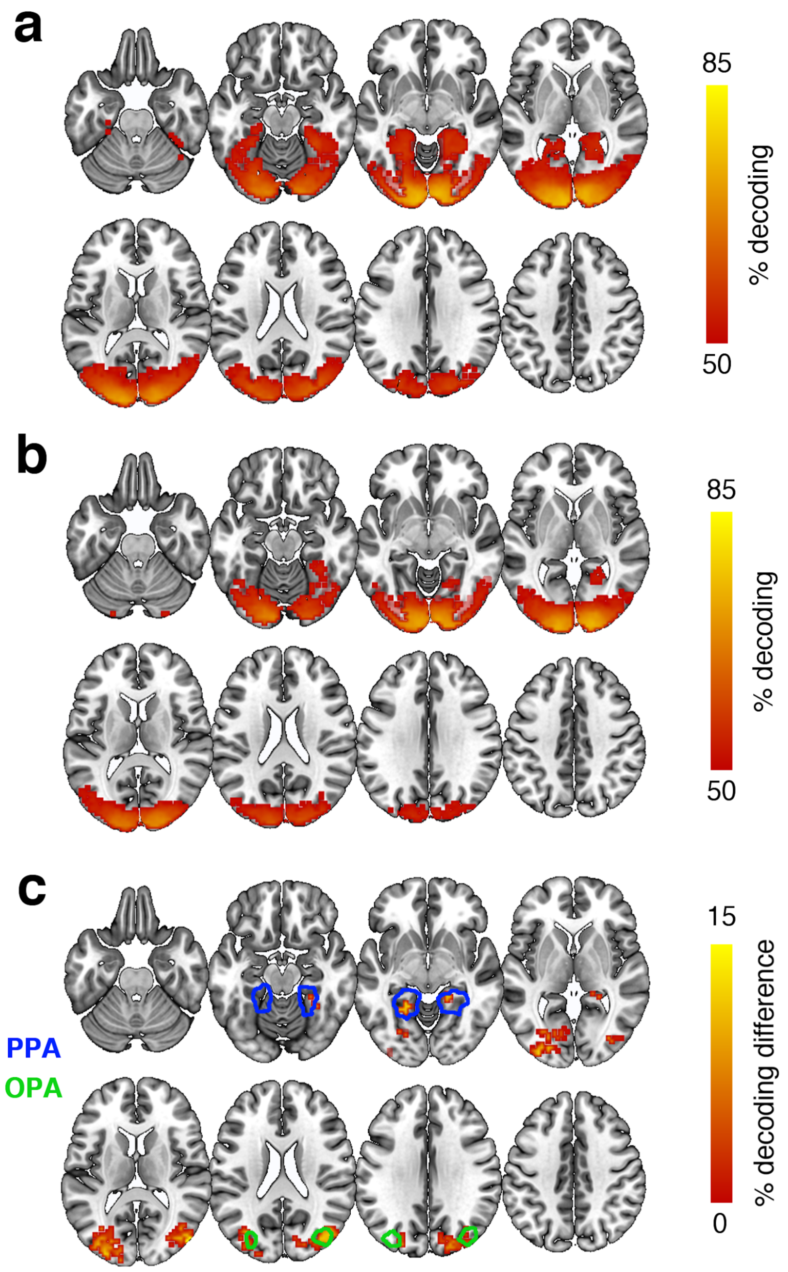


*Figure S2.* MVPA searchlight results in occipitotemporal cortex. Spatially intact and spatially jumbled scenes were discriminable in widespread regions of the visual cortex, both when presented upright (a) and inverted (b). Critically, when subtracting decoding in the upright and inverted conditions, we found inversion effects in regions overlapping with the typical locations of OPA and PPA (masks taken from Julian et al., 2012) (c), which indicated genuine sensitivity to spatial scene structure. Only voxels exhibiting significant effects ($p_{corr}$<.05) are shown.

The searchlight analysis yielded widespread significant decoding between spatially intact and jumbled upright scenes, covering early and high-level visual cortex (Figure S2a). For inverted scenes, this decoding was less pronounced (Figure S2b). Critically, we found inversion effects (i.e., better decoding for the upright, compared to the inverted scenes) in areas around the transverse occipital sulcus, corresponding to the location of OPA and areas around the parahippocampal cortex, corresponding to the location of PPA (Figure S2c). No significant inversion effects were found for categorical scene structure.

These analyses strongly support the results of our ROI-based analysis (Figure 3b/e), which revealed genuine sensitivity to spatial scene structure in the OPA and PPA.

**fMRI decoding – additional ROIs**

In addition to the scene-selective OPA and PPA, we also performed MVPA on responses in scene-selective retrosplenial cortex (RSC) and object-selective lateral occipital cortex (LO). These ROIs were defined similarly to OPA and PPA: For both regions, we used a functional template mask (Julian et al., 2012), and within this mask defined the voxel exhibiting the greatest *t*-value in a scene>object (RSC) or an object>scrambled (LO) contrast. Then, the ROIs were constructed as 125-voxel spheres around this peak voxel, and concatenated for the left and right hemispheres. After extracting responses from these ROIs, we performed the same decoding analyses (Figure S3a/c) as for the other ROIs.

For RSC, we did not find significant decoding between the spatially intact and spatially jumbled scenes (Figure S3b), neither in the upright, $t(19)=2.49$, $p_{corr}=.066$, nor the inverted condition, $t(19)=0.13$, $p_{corr}>1$. No significant inversion effect was observed, $t(16)=1.82$, $p_{corr}=.26$. Similarly, no significant effects were found when decoding between categorically intact and categorically jumbled scenes (Figure S3d), all $t<1.64$, $p_{corr}>.35$. These results suggest that scene structure is not represented in RSC.

For LO, we found significant decoding between spatially intact and spatially jumbled scenes (Figure S3b), both in the upright, $t(19)=4.19$, $p_{corr}=.001$, and in the inverted condition, $t(19)=3.48$, $p_{corr}=0.008$. However, no inversion effect was found, $t(16)=0.47$, $p_{corr}>1$. No significant effects were found when decoding between categorically intact and categorically jumbled scenes (Figure S3d), all $t<2.19$, $p_{corr}>.12$. These results suggest that only scene-selective regions, but not object-selective regions of the occipital cortex are genuinely sensitive to spatial scene structure.

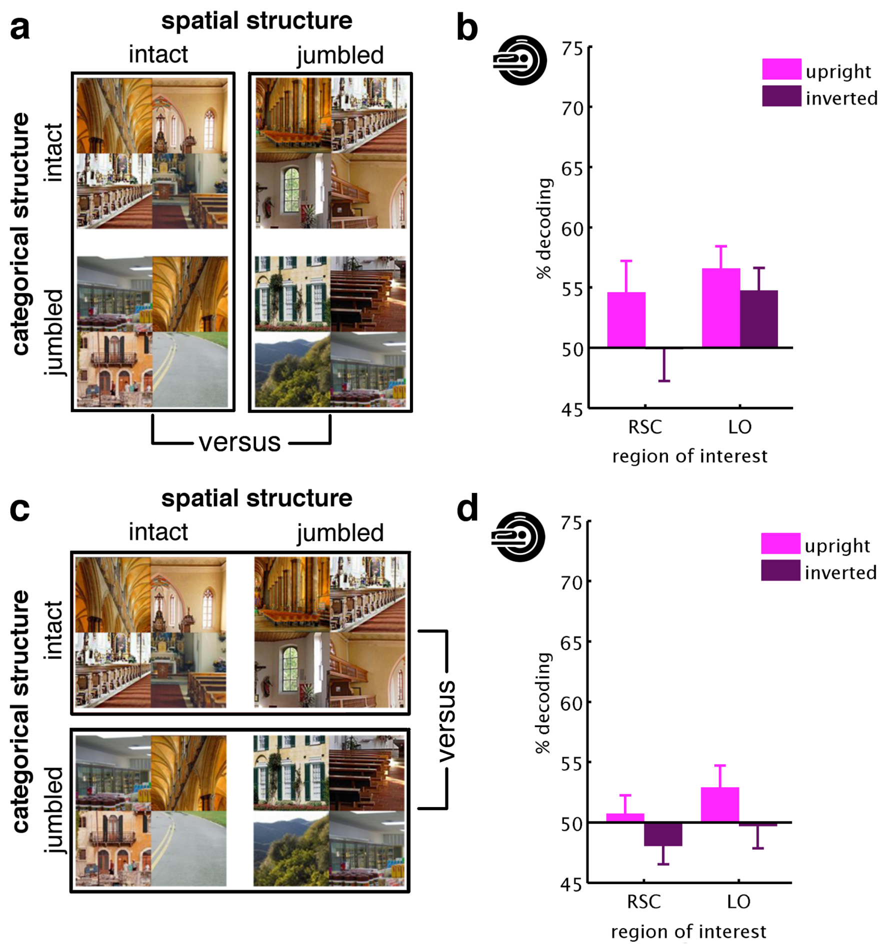


*Figure S3.* MVPA results in RSC and LO. To reveal sensitivity to spatial scene structure, we decoded between scenes with spatially intact and spatially jumbled parts (a). Scene-selective RSC did not show any significant decoding of spatial scene structure and no inversion effects. Object-selective LO showed significant decoding between spatially intact and spatially jumbled scenes, but no significant inversion effect (b). To reveal sensitivity to categorical scene structure, we decoded between scenes with categorically intact and categorically jumbled parts (c). In this analysis, no significant decoding and no inversion effects were found for both regions (d).

**fMRI decoding – varying voxel counts in V1**

To explore whether the results in V1 changed as a function of ROI size, we selected different numbers of voxels, depending on their probability to belong to V1, taken from the Wang et al. (2015) atlas. The resulting V1 sizes varied between 1032 (10% probability cutoff) and 87 voxels (60% probability cutoff).

For each of these voxel counts, we re-performed the main decoding analysis, where we decoded between (1) spatially intact and spatially jumbled scenes and (2) categorically intact and categorically scrambled scenes (Figure S4a/c).

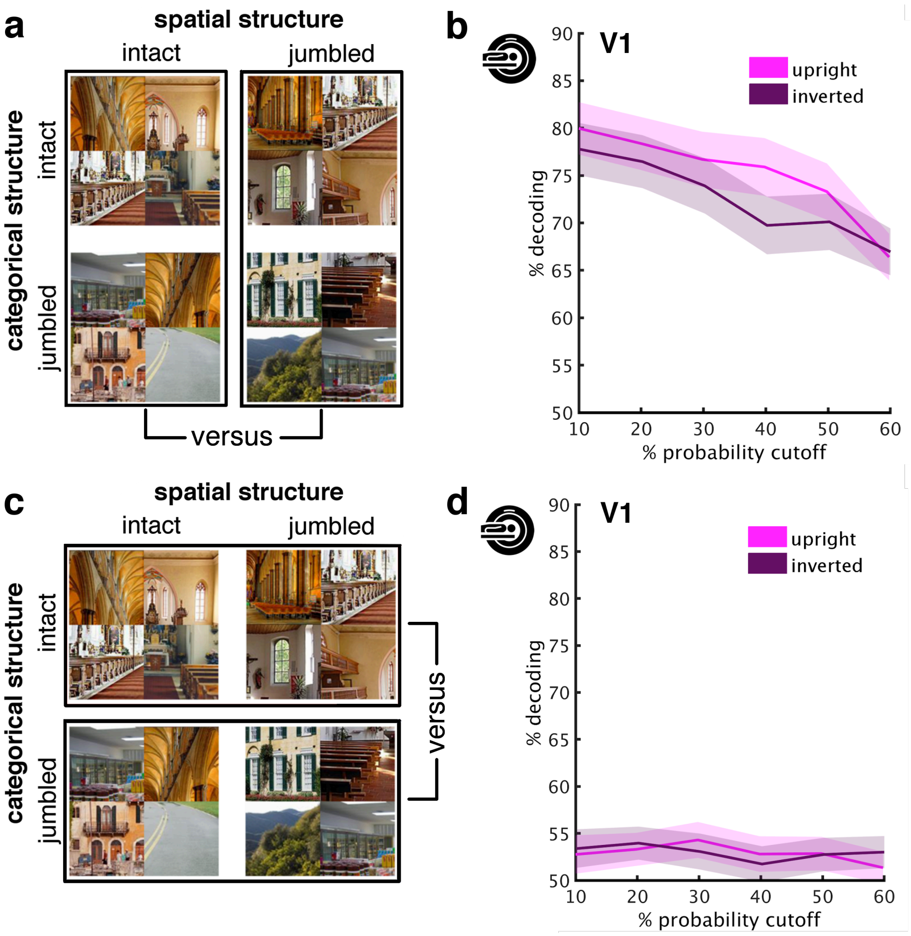


*Figure S4.* Results for different voxel counts in V1. Results were highly similar across different V1 sizes, ranging from voxels belonging to V1 with probabilities ≥10% (1032 voxels) to voxels belonging to V1 with probabilities ≥60% (87 voxels). This shows that independently of the region's size, there is no reliable sensitivity to scene structure (i.e., no robust inversion effects) in early visual cortex. Error margins reflect standard errors of the difference.

Independently of the voxel counts, we found similar results as in the main analysis (Figure 3b/e). Spatial structure could be decoded reliably from V1 activations, both in the upright and inverted conditions, all $t(19)>7.32$, $p_{corr}<.001$ (Figure S4b). We observed an inversion effect only for the 40% probability cutoff, $t(16)=3.01$, $p_{corr}=.025$, but not all other cutoffs, all $t(16)<1.62$, $p_{corr}>.37$. Across the different voxel counts, we did not find a difference between the upright and inverted conditions, $F(1,16)=2.19$, $p_{corr}=.47$, suggesting no genuine inversion effects in V1. Similarly, we did not observe any significant inversion effects when looking at categorical scene structure, all $t(16)<1.07$, $p_{corr}>.90$ (Figure S4d). These results corroborate our finding that V1 does not exhibit robust sensitivity to scene structure.

**fMRI decoding – varying voxel counts in OPA / PPA**

To explore whether the results in OPA and PPA changed as a function of ROI size, we selected different numbers of voxels by varying the number of voxels selected around the localizer peak activation of each hemisphere (see Materials and Methods). Each ROI's size was varied between 25 voxels and 225 voxels for each hemisphere (i.e., 50 to 450 voxels for the collapsed ROI).

For each of these voxel counts, we re-performed the main decoding analysis, where we decoded between (1) spatially intact and spatially jumbled scenes and (2) categorically intact and categorically scrambled scenes (Figure S5a/c).

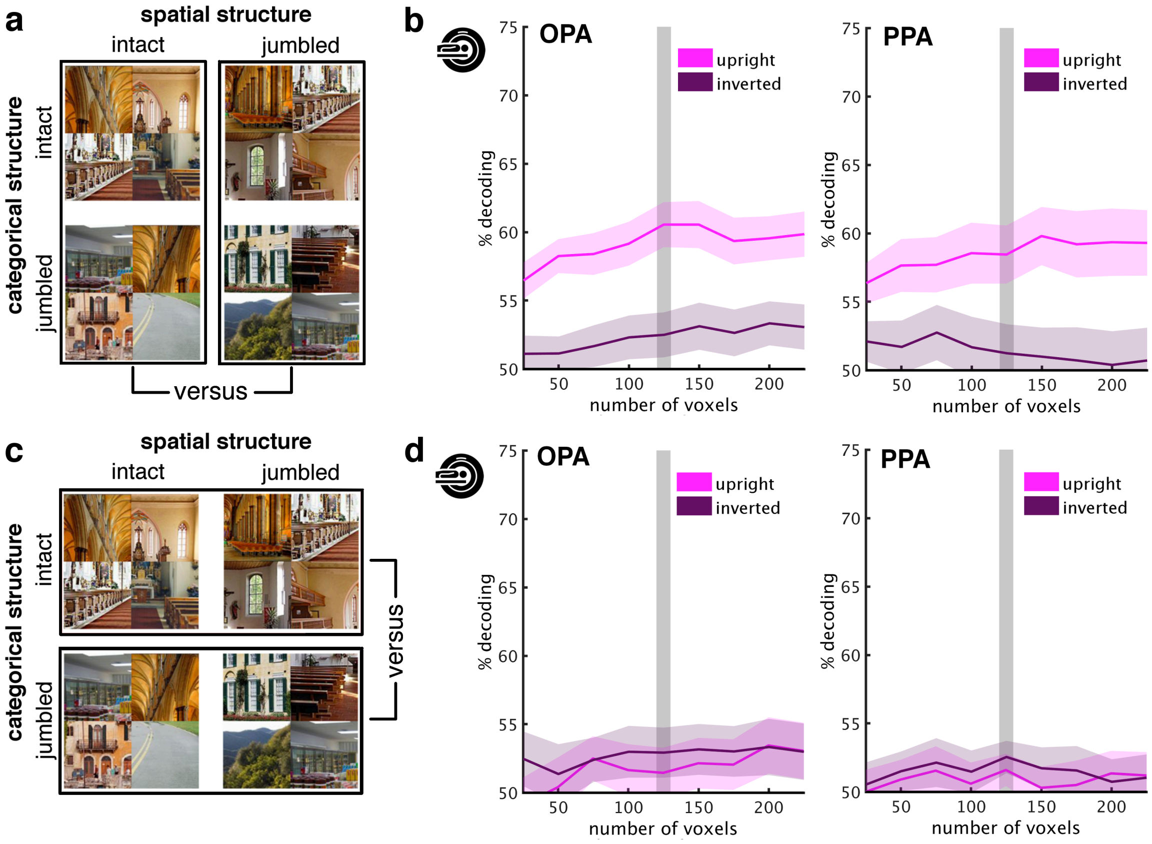


*Figure S5.* To explore whether the results in scene-selective cortex change as a function of ROI size, we selected the nearest 25 to 225 voxels around the individual participants' localizer peaks. Critically, the results were highly similar across the different ROI sizes, showing that the observed effect of sensitivity to spatial structure in scene-selective cortex is not strongly dependent on the number of voxels considered part of each ROI. Error margins reflect standard errors of the difference. Shaded gray bars mark the 125-voxel spheres used in the main analysis.

We found that results were largely independent of region size. Spatial structure could be reliably decoded for upright scenes in OPA, all $t(19)>3.70$, $p_{corr}<.005$, and PPA, all $t(19)>5.19$, $p_{corr}<.003$ (Figure S5b). This decoding was significantly weaker for in the inverted condition, both in OPA, all $t(16)>3.37$, $p_{corr}<.012$, and PPA, all $t(16)>2.50$, $p_{corr}<.071$, indicating inversion effects across all voxels counts. By contrast, we did not observe any significant inversion effects when looking at categorical scene structure, neither in OPA, all $t(16)<0.61$, $p_{corr}>1$, nor in PPA, all $t(16)<0.51$, $p_{corr}>1$ (Figure S5d). These results show that our finding of robust sensitivity to spatial scene structure in scene-selective cortex cannot be attributed to the ROI definition criteria applied.

**EEG decoding – classifying scene category**

Our data suggest that categorically intact and categorically shuffled scenes were not represented differently during the experiments. Could this result be explained by an absence of category information from neural signals in the first place?

To investigate how well cortical representations tracked the scenes' categorical content, we performed a decoding analysis on the EEG data in which we classified scenes into the four categories used in the experiment (church, house, supermarket, street). Note that this analysis could not be performed for the fMRI experiment, where scenes of all categories were intermixed within each block of the block design.

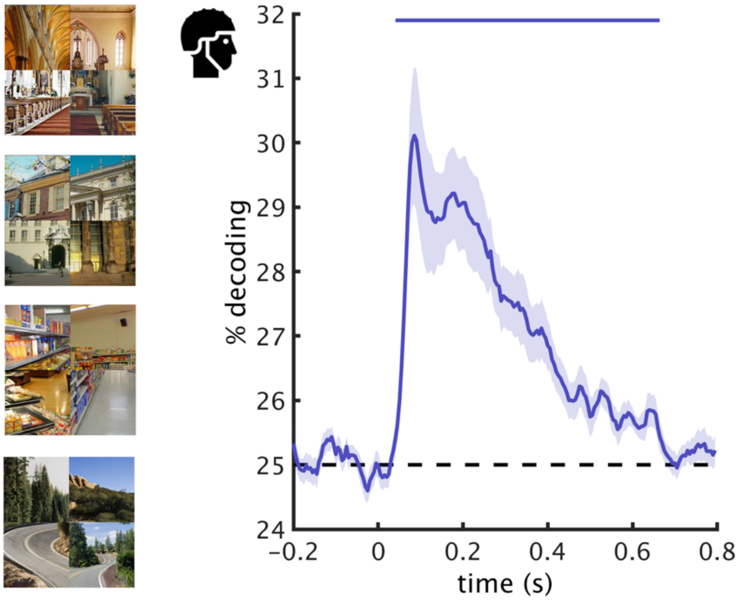


*Figure S6.* Decoding of scene categories from EEG signals. The four scene categories (example stimuli on the left) could be reliably decoded from the EEG data between 45ms and 660ms, showing that the neural data contained robust information about scene category. Error margins reflect standard errors of the mean. Significance markers denote above-chance decoding ($p_{corr}$<.05).

For decoding scene category from the EEG signals, we only used the conditions where scene category remained intact across the four scene parts. For each of these four conditions separately, we then performed a four-way decoding analysis in a leave-one-trial-out fashion (see Materials and Methods for details on the decoding procedure), and subsequently averaged across these analyses. Note that the purpose of this analysis was to show that the EEG signals contained reliable category information. Further analyses on the nature of this category information are beyond the scope of the current paper.

Across the conditions analyzed, we found that scene category information was robustly decodable between 45ms and 660ms after scene onset, peak $z>3.71$, $p_{corr}<.001$ (Figure S6). This shows that there was robust category information in the EEG signals, although across scenes there was no sensitivity to the scenes' categorical structure (i.e., whether categorical content matched within a scene or not).
